# Framework for reanalysis of publicly available Affymetrix^®^ GeneChip^®^ data sets based on functional regions of interest

**DOI:** 10.1101/126573

**Authors:** Ernur Saka, Benjamin J. Harrison, Kirk West, Jeffrey C. Petruska, Eric C. Rouchka

**Author notes:** Corresponding author: Eric C. Rouchka, Department of Computer Engineering and Computer Science University of Louisville, Louisville KY, USA. Email addresses: ES BJH KW JCP ECR.

## Abstract

**Background:** Since the introduction of microarrays in 1995, researchers world-wide have used both commercial and custom-designed microarrays for understanding differential expression of transcribed genes. Public databases such as ArrayExpress and the Gene Expression Omnibus (GEO) have made millions of samples readily available. One main drawback to microarray data analysis involves the selection of probes to represent a specific transcript of interest, particularly in light of the fact that transcript-specific knowledge (notably alternative splicing) is dynamic in nature.

**Results:** We therefore developed a framework for reannotating and reassigning probe groups for Affymetrix^®^ GeneChip^®^ technology based on functional regions of interest. This framework addresses three issues of Affymetrix^®^ GeneChip^®^ data analyses: removing nonspecific probes, updating probe target mapping based on the latest genome knowledge and grouping probes into gene, transcript and region-based (UTR, individual exon, CDS) probe sets. Updated gene and transcript probe sets provide more specific analysis results based on current genomic and transcriptomic knowledge. The framework selects unique probes, aligns them to gene annotations and generates a custom Chip Description File (CDF). The analysis reveals only 87% of the Affymetrix^®^ GeneChip^®^ HG-U133 Plus 2 probes uniquely align to the current hg38 human assembly without mismatches. We also tested new mappings on the publicly available data series using rat and human data from GSE48611 and GSE72551 obtained from GEO, and illustrate that functional grouping allows for the subtle detection of regions of interest likely to have phenotypical consequences.

**Conclusion:** Through reanalysis of the publicly available data series GSE48611 and GSE72551, we profiled the contribution of UTR and CDS regions to the gene expression levels globally. The comparison between region and gene based results indicated that the detected expressed genes by gene-based and region-based CDFs show high consistency and regions based results allows us to detection of changes in transcript formation.

## Background

A DNA microarray (DNA chip or biochip) is a technology used to identify and measure the expression level of specific mRNA molecules in order to ascertain transcriptional profiles in response to differing conditions. The most commonly used microarray is the Affymetrix^®^ GeneChip^®^ family of arrays. Each GeneChip^®^ consists of a silicon chip with fixed locations called cells, spots or features [1]. Each spot contains millions of identical 25 base oligonucleotides (probes) which are selected to be complementary to various transcript regions of a gene [2]. In order to determine transcript expression, which directly infers gene expression, groups of 11-20 probes matching the same gene/transcript are arranged in a probe set. Given a particular Affymetrix^®^ GeneChip^®^ platform, the design of the probes is fixed based on earlier genome assemblies and annotation available at that time. Since the design of the first Affymetrix^®^ GeneChip^®^, rapid progress has been made in genome sequencing resulting in more accurate databases of annotated coding and noncoding genes.

The significant differences between old and new genome assemblies and annotations make it necessary to update probe-gene targeting according to current knowledge to get more accurate interpretations from experimental results. Affymetrix^®^ does attempt to provide compatibility between genomic changes by updating links between probe sets and their corresponding genes/transcripts via NetAffx™ [3]. Table 1 shows release dates of source databases used by Affymetrix^®^ for both the incorporated version and the most recently available version. In all cases, there is at least a two year difference between the incorporated and most recent release dates which can lead to inconsistent interpretation.

**Table 1.**
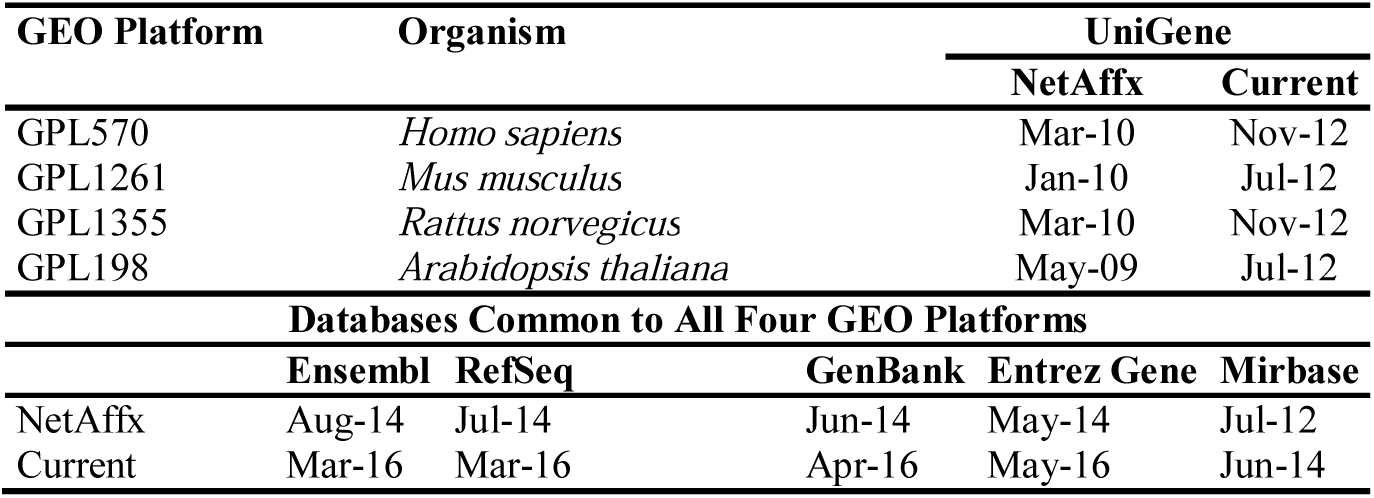
Release dates of databases used by NetAffx v35 annotations and current database versions.

| GEO Platform | Organism | UniGene |  |  |  |
| --- | --- | --- | --- | --- | --- |
|  |  | NetAffx | Current |  |  |
| GPL570 | *Homo sapiens* | Mar-10 | Nov-12 |  |  |
| GPL1261 | *Mus musculus* | Jan-10 | Jul-12 |  |  |
| GPL1355 | *Rattus norvegicus* | Mar-10 | Nov-12 |  |  |
| GPL198 | *Arabidopsis thaliana* | May-09 | Jul-12 |  |  |
| Databases Common to All Four GEO Platforms |  |  |  |  |  |
|  | Ensembl | RefSeq | GenBank | Entrez Gene | Mirbase |
| NetAffx | Aug-14 | Jul-14 | Jun-14 | May-14 | Jul-12 |
| Current | Mar-16 | Mar-16 | Apr-16 | May-16 | Jun-14 |

In addition, updating links between probe sets and their corresponding genes/transcripts does not provide a solution for problems caused by individual probes such as single nucleotide polymorphisms (SNPs) [4, 5], probes that target genes other than the designated gene of a probe set, and probes that no longer align to a genomic location. For example in the Affymetrix^®^ GeneChip^®^ HG-133 Plus 2 array, a total of 40,680 probes out of 603,158 (excluding quality control probes) do not have a perfect match to the most recent human genome assembly (hg38).

Even though the design of the probes is fixed, the methods with which the resulting experiments can be analyzed are dynamic in nature due to the ability to annotate and arrange probes into uniquely defined groupings. This is particularly important since there are publicly available repositories of microarray datasets, such as NCBI’s Gene Expression Omnibus (GEO) [6] which contains 1,802,922 different samples as of 5/18/2016 that can be reanalyzed computationally based on current knowledge without the need for new biological experiments. As a case in point, each of the four most commonly used species have samples that have been analyzed using the original CDFs (Table 2).

**Table 2.**
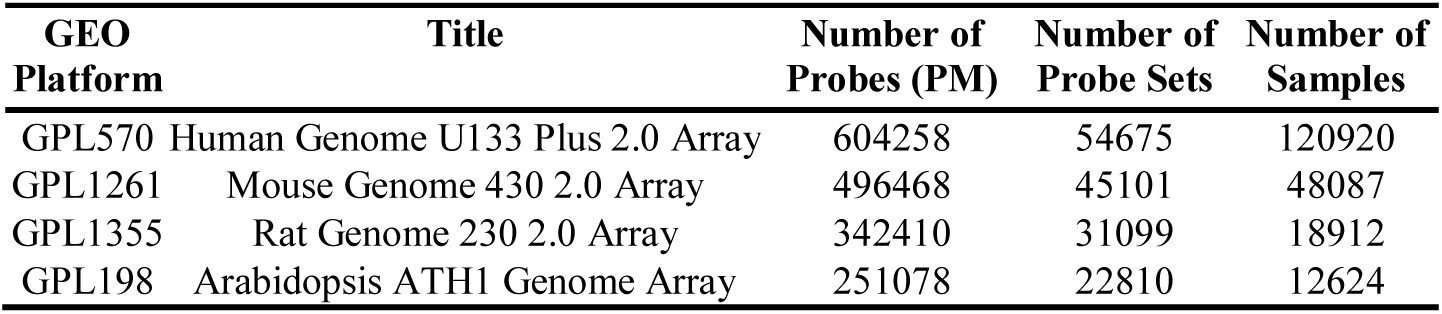
Top Affymetrix^®^ *in situ* oligonucleotide arrays found in GEO.

| GEO Platform | Title | Number of Probes (PM) | Number of Probe Sets | Number of Samples |
| --- | --- | --- | --- | --- |
| GPL570 | Human Genome U133 Plus 2.0 Array | 604258 | 54675 | 120920 |
| GPL1261 | Mouse Genome 430 2.0 Array | 496468 | 45101 | 48087 |
| GPL1355 | Rat Genome 230 2.0 Array | 342410 | 31099 | 18912 |
| GPL198 | Arabidopsis ATH1 Genome Array | 251078 | 22810 | 12624 |

Several research groups have reassigned probes into new probe sets by creating their own custom Chip Description Files (CDF) [7-13], which are specially formatted files used to store the layout information for an Affymetrix^®^ GeneChip^®^ array. Given a CDF, the intensity values of probes located in the CEL file can be extracted and summarized as a defined probe set to detect the expression level of genes or transcripts. These approaches have a similar workflow of mapping probes but differ in terms of the groupings of probe sets, including: data source used, the selected target level (gene or transcript), whether to create probe sets from scratch or redesign the existing groups and sharing probes between probe sets.

In terms of annotations used, most approaches have mapped the probe sequences to the transcripts obtained from one or more databases such as GenBank, NCBI RefSeq and Ensembl. Unlike other approaches, Harbig et al. [14] mapped to the target sequences of probes obtained from Affymetrix^®^ rather than the actual mRNA sequences themselves, where the target sequence is an exemplar region of a specific transcript ≤ 600 bases in length. After mapping, they grouped probes to unique transcripts or genes based on the mapping results. Some approaches update the original probe set groups by removing select probes and changing the link between probe set and gene/transcript. The most comprehensive study for probe annotation remapping was achieved by Dai et al (brainarray CDFs) [8]. Rather than focusing on one reference database or combining multiple sources to create one custom CDF, they mapped probes to different annotation databases and created a specific custom CDF for each database.

Although the inherent effects of using dated probe gene mapping designs to analyze microarray data sets might seem obvious, the overwhelming majority of experimental results have only been analyzed using the original CDFs designed by Affymetrix^®^. For example as of May 2016, GEO has 120,920 samples which were analyzed via the original Affymetrix^®^ CDFs for the HG-U133 Plus 2 array (Table 2). On the other hand only 6,403 samples were analyzed using custom CDFs, mostly produced by brainarray (Table 3). Given that fewer than 5% of all samples in GEO have been analyzed by alternative CDFs, an opportunity exists to reanalyze existing datasets according to updated transcript knowledge or functional regions of interest.

**Table 3.** Alternative CDFs for the top Affymetrix^®^ *in situ* oligonucleotide arrays found in GEO.

| GEO Platform | Number of Alternative CDFs | Number and Percent of Samples Using Alternative CDFs |
| --- | --- | --- |
| GPL570 | 54 | 6403 (5.0%) |
| GPL1261 | 36 | 1984 (4.0%) |
| GPL1355 | 12 | 460 (2.4%) |
| GPL198 | 9 | 642 (4.8%) |

While microarrays have been successfully utilized for understanding differential expression at the gene or probe set level, less attention has been given to the potential analysis at the individual exon, alternative transcript, and untranslated region (UTR) level. While the selection bias of probes on the 3’ ends of genes for earlier iterations of Affymetrix^®^ GeneChip^®^ designs presents limitations on the completeness of transcript information, more recent designs allow for a more complete coverage of exons and exon junctions. However, information concerning individual exons can still be extracted from earlier GeneChip^®^ designs, particularly in the 3’ UTR regions that have been shown to play important roles in cancer [15-17], development [18-22], and localization in the nervous system [23-27]. In fact, over 40% of genes have been shown to generate multiple mRNAs with variable 3’ UTR lengths [28]. These 3’ UTRs harbor binding sites for molecules including microRNAs (miRNAs) and RNA-binding proteins. Thus, mRNA isoforms with lengthened 3’ UTRs have increased numbers of sites for these cis-interacting factors. The diversity of 3’ UTRs is predominantly regulated by alternative polyadenylation (APA), which employs alternative mRNA cleavage sites that lie progressively distal to the stop codon. APA-driven mRNA diversity is required for normal physiology, and misregulation of this process is associated with diverse disease states [29]. We therefore have developed a framework for analysis of Affymetrix^®^ GeneChip^®^ data by regrouping probes into probe sets based on Ensembl annotations at the gene, transcript, individual exon, and UTR levels in order to detect changes in gene expression that may occur within specific regions of the transcript.

## Methods

We developed an Affymetrix^®^ GeneChip^®^ probe remapping protocol at the level of genes, transcripts, untranslated regions (UTRs), coding sequences (CDS) and individual exons based on the latest genome (hg38, mm10, rn6) and Ensembl annotations (ENS-85) for human, mouse, and rat. The protocol takes annotations in a General/Gene Transfer Format (GTF) [30] file, generates a custom CDF where probes are grouped into probe sets based on region (UTR, CDS, individual exon), transcript or gene level. Here, we define individual exons as coding exons within protein coding genes, or all exons within structural RNAs (such as miRNA and lncRNA). In effect, the individual exons refer to all non-UTR portions of exons. Fig. 1 shows the flow chart of annotation and grouping of probes based on the region of a gene. It is composed of three main steps: mapping probes to the genome, annotation of probes, and assignment of probes to probe sets based on annotations.

**Fig. 1.**
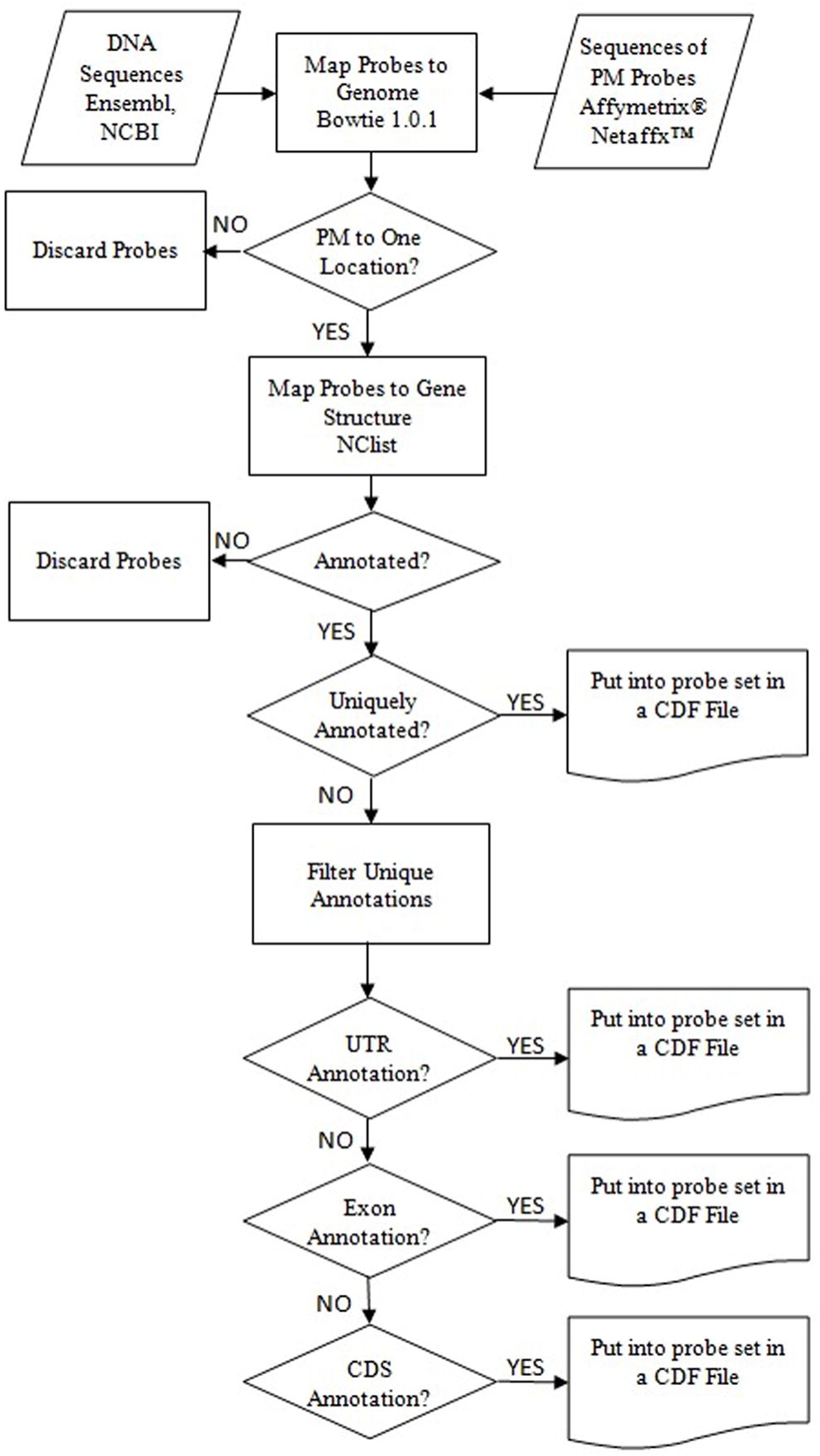
Flow chart for region-based probe annotation framework.

### Mapping of perfect match probes to a genome

PM probe sequences, which can be obtained from the Affymetrix^®^ Netaffx™ web site, are aligned to the indexed genome using Bowtie version 1.0.1 [31] with the parameters -v 0 and −m 1, requiring that probes align to a single genomic location with 100% identity, thereby reducing cross-hybridization effects. Note that Bowtie version 1 is best at aligning shorter sequences (25-50 bp) as found with microarray probes while the most recent versions of Bowtie are optimized for long sequence reads (>50 bp). Mismatch (MM) probes are not considered in the mapping step, although they could theoretically map uniquely to genomic regions. Rather, the MM probes are set aside and are included with their corresponding PM probe during the final CDF construction step once the PM probes have been assigned to a probe set. During this analysis, only probes perfectly matching to a region are considered. Therefore, probes crossing splice junctions will be discarded.

### Annotation of perfect match probes via nested containment list (NCList)

Probes are annotated based on the overlap between probes and genomic intervals by the following steps.

I. GTF [30] files for the mouse, rat, and human genome were obtained from the Ensembl ftp server [32]. Each GTF is a tab-delimited text file used to represent gene structure information, including the start and end positions of a gene together with chromosome location. Each structure is tagged with a feature which can be gene, transcript, exon, start_codon, stop_codon, CDS or UTR. Ensembl GTFs were used since the annotations are determined by an automated system based on experimentally verified data combined from multiple databases such as RefSeq, EMBL and UniProtKB. It also contains manual curation for selected species.
II. A nested containment list (NCList) [33] was created for each chromosome from intervals (start and end points) of gene structures. The intervals of the NCList were selected based on the target of the probe sets. When the probe sets were constructed based on regions of a gene, we used UTR, individual exon and CDS intervals. For gene/transcript targeted probe sets, we used gene/transcript intervals.
III. Probe intervals were searched in the NCList and annotated according to the overlapping results. Probes were split based on the matched chromosome. Each probe group interval was searched in the same chromosome’s NCList. When an overlap was found, the probe was annotated with the list node. Only complete overlaps were accepted; both the low and high ends of the interval have to be included in the list node. The probes which did not overlap the nodes were discarded. As a result, probes partially overlapping UTRs, individual exons, and CDS regions will not be included at the region and gene level, but will be present at the transcript level.
IV. A probe’s start and end points may overlap multiple gene structures. It may overlap with the UTR and exon region of the same gene or with multiple genes or transcripts. In order to remove cross hybridization and ensure probes uniquely map to a single region, gene or transcript, we choose one of the annotations for each probe and remove the remaining matches. The rule for assigning these probes occurs with the following priority (I) 5’ and 3’ UTRs; (II) exons; (III) CDS. Thus, although UTR regions technically occur within exons, the more specific UTR assignment will be used. When the annotation was based on gene or transcript the first obtained annotation was selected.
V. Probes with the same annotation were grouped together to form a probe set. Fig. 2 shows the grouping of probes for three types of CDFs. These CDFs are:

- Region-based CDF: Probe sets are designed to target a specific region of a gene and consist of probes which map to the same region (UTR, individual exon, CDS) of a gene. In Fig. 2, green probes were mapped to the UTR region of Gene_1; therefore, those probes cluster together to form the Gene_1 UTR region probe set. Based on the same logic, blue colored probes form the probe set for Gene_1 exon and pink colored probes form the probe set for Gene_1 CDS.
- Gene-based CDF: Probe sets are designed to target genes and consist of probes which map to the same gene. In Fig. 2, green, blue and pink colored probes, which mapped to Gene_1, cluster together to form the Gene_1 probe set.
- Transcript-based CDF: Probes that map to same transcript of a gene compose a probe set. In Fig. 2, the orange and red arrow show the start and end positions of Transcript_1 and Transcript_2. The probes mapped to the Transcript_1 (two greens, two blue and two pink) cluster together to form the probe set for Transcript_1.
VI. Probe sets were saved into binary and ASCII format CDF files. The CDF files were created via the affxparser [34] Bioconductor package. In addition to the probes specific for a particular gene, Affymetrix^®^ GeneChips^®^ contain a number of different control probes such as probes that are added during sample preparation, providing evidence that assay was performed properly. We added those probe sets to our CDFs without any change. R CDF libraries were created via the makecdfenv [35] R Bioconductor package. The custom CDFs for three species (rat, mouse, and human) can be obtained from bioinformatics.louisville.edu/RegionCDFDesc.html

**Fig. 2.**
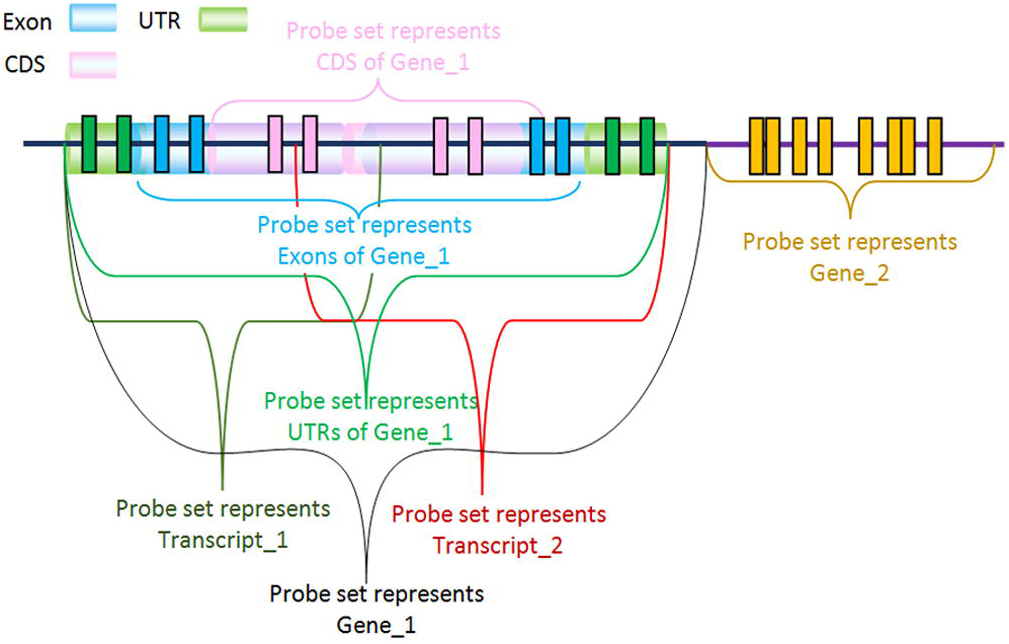
Creating probe sets for different types of custom CDF based on probe mapping to gene regions.

### Probe Set Naming

Since GTF files obtained from Ensembl were used, Ensembl gene ids were employed to distinguish different genes and Ensembl transcript ids were used to distinguish different transcripts. When the generated CDF was based on regions of genes, the region was suffixed to the Ensembl gene id. Table 4 shows example probe set names taken from custom CDFs for the Affymetrix^®^ GeneChip^®^ HG-133 Plus 2.

**Table 4.** Custom CDF naming examples.

| CDF Type | Probe Set Name |
| --- | --- |
| Region-based | ENSG00000001036_exon_- |
|  | ENSG00000001084_UTR_- |
|  | ENSG00000001167_CDS_+ |
| Gene-based | ENSG00000001461 |
| Transcript-based | ENST00000489806 |

We applied our framework to the three most widely used GeneChips^®^: HG-U133 Plus 2, Rat 230 2.0 and Mouse 430 2.0 (summarized in Tables 5 and 6). We also examined the effect of probe reannotation over the differentially expressed genes. Three types of CDFs were created for every selected organism. Our results discussed here are restricted to the analysis of the HG-U133 Plus 2 and Rat Genome 230 2.0 GeneChip^®^ for brevity. After CDF creation, we reanalyzed the publicly available data series GSE48611 [36] and GSE72551 [37] from GEO via our custom CDFs.

**Table 5.**
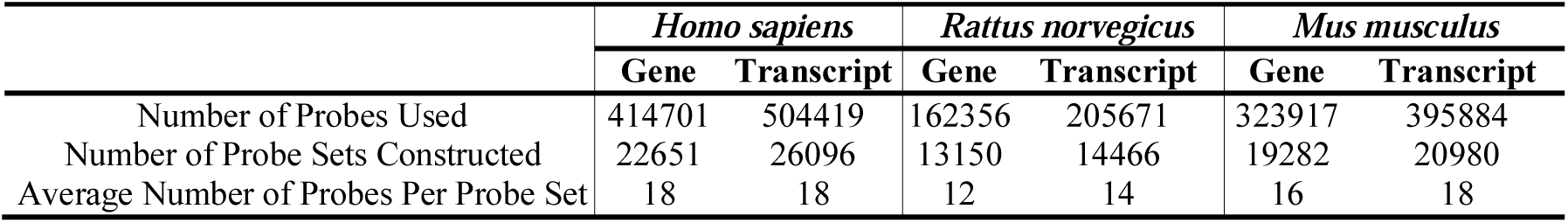
Summary of probes used for custom gene and transcript based Custom CDFs.

**Table 6.** Summary of probes used for region based custom CDFs.

|  | *Homo sapiens* | *Rattus norvegicus* | *Mus musculus* |
| --- | --- | --- | --- |
| Number of Probes Aligned to Genome | 822681 | 321905 | 637942 |
| Number of Probes Used | 414701 | 162356 | 323917 |
| Number of Probe Sets Constructed | 33916 | 19839 | 28963 |
| Average Number of Probes Per Probe Set | 12 | 8 | 11 |

## Results

### Custom CDF Generation

#### Probes Mapping to the Genome

Using the bowtie parameters as discussed in the methods section, we were able to identify probes that uniquely map with 100% identity for each of the respective genomes. As a result, 87% PM probes of the HG-U133 Plus 2, 84% PM probes of the Rat 230 2.0 and 86% PM probes of the Mouse 430 2.0 were uniquely mapped to the genome and were used in the subsequent steps (Table 7).

**Table 7.** Number of mapped probes for custom CDF construction.

| GeneChip® | Number of PM Probes | Number of PM Probes Mapped Uniquely | Number of PM Probes Mapped to Multiple Locations | Number of PM Probes Not Aligned |
| --- | --- | --- | --- | --- |
| Human Genome U133 Plus 2.0 Array | 603158 | 525985 | 36493 | 40680 |
| Rat Genome 230 2.0 Array | 341459 | 288319 | 26027 | 27113 |
| Mouse Genome 430 2.0 Array | 495374 | 427758 | 28444 | 39173 |

#### Probe Annotations and Probe Sets

To annotate probes, we mapped uniquely aligned probes to gene regions using the most recent Ensembl genome and GTF file for each respective organism. We used the specific regions based on the custom CDF type (gene, transcript or region-based). Consequently we produced three types of custom CDFs (Tables 5 and 6).

The human gene based CDF has 22,651 custom designed probe sets composed from 414,701 probes and 62 original control probe sets. 442,025 annotations were identified between genes and the probes. 27,324 annotations were filtered after shared probes were removed. In order to validate our probe set annotations, we compared the original CDF probe sets with the custom CDF. A total of 21,585 annotated genes were shared between the two CDFs, with 3,068 unique to the original CDF, and 1,066 unique to our custom CDF. In order to determine why some genes were not covered in our CDF, we examined those unique to the original CDF. First we obtained the probe sets which represent these genes in the original CDF, yielding 2,781 probe sets. We retrieved both the PM and MM probe sequences for each of these. We observed that for 667 probe sets, every probe was removed during probe mapping to the genome due to either non-unique mappings or mapping rates less than 100%. 30,150 probes from the remaining 2,114 probe sets were not used in our CDF since they either did not map to the genome or they were MM probes. 14,028 probes were used in our newly constructed probe sets which target different genes than the original assignment by Affymetrix^®^ and 2,656 probes were not aligned to gene structures and not annotated. As a result, the differences between the original CDF and our method occurs because of probes removed during genome alignment, probes that no longer map to gene structure or probes that map to gene structures different from the original annotation.

For the rat 230 2.0 GeneChip^®^, the restriction of three probes per probeset yields 12,534 uniquely identified Ensembl genes at the gene level. We determined that for this specific GeneChip^®^, reorganization of the Affymetrix^®^ probes into mRNA region-specific probesets provides 4,024 unique Ensembl gene identifiers with probesets in both the 3’ UTR and CDS. Using this subset of probesets, differential expression of the CDS can then be compared to the 3’ UTR.

#### Analysis with Custom CDFs

We reanalyzed the publicly available data series GSE72551 and GSE48611. Both of these studies involve the nervous system, where differences in 3’ UTRs are likely to have phenotypic effects on transcript localization. The GSE72551 data series examines gene expression changes associated with collateral sprouting and includes 5 naïve controls, 7 replicates at day 7 post-surgery and 7 replicates at day 14 postsurgery. The GSE48611 data series examines Down syndrome gene expression monitoring. This data set includes mRNA samples from the isogenic trisomy of chromosome 21 (Ts21) and control pluripotent stem cells (iPSCs) (DS1, DS4, and DS2U) between passages 24 and 48 and from day 30 neurons. Three biological replicates were present for each condition. Prior to analysis, we removed probe sets with two or fewer probes from the custom CDFs in order to achieve more accurate results for target expression levels. Robust Multiarray Averaging (RMA) normalization [38] was used for preprocessing. A p-value 0.05 was used as the threshold for all experiments.

In the GSE72551 data series, differentially expressed genes (DEGs) were determined for two pairwise comparisons: naïve vs. both 7 and 14 days using region and gene based custom CDFs. We also reanalyzed the data using brainarray Ensembl CDF version 20. Fig. 3 shows a Venn diagram representing the number of differentially expressed genes using region, gene and brainarray custom CDFs for both cases.

**Fig. 3.**
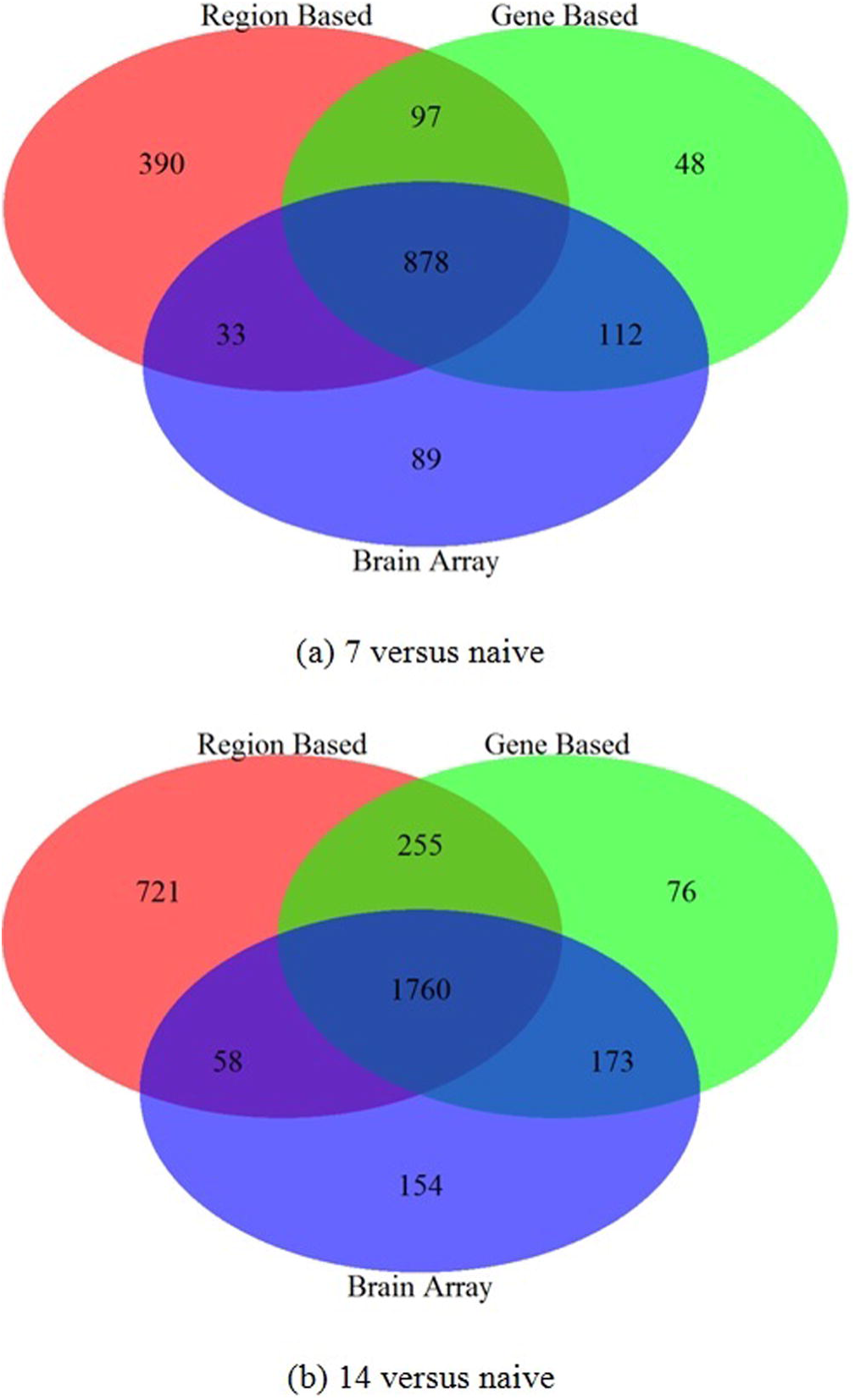
Number of common and different differentially expressed genes using region, gene and brain array custom CDFs. **a** Day 7 versus naïve. **b** Day 14 versus naïve.

Further examination of the 7 day versus naïve ENSEMBL genes found to be differentially expressed in either the gene-based or region-based CDF shows high concordance, with 975 ENSEMBL genes determined to be differentially expressed using both CDFs (Fig. 3a). Examination of the p-values shows a significant correlation between both the gene and the 3’ UTR region (r=0.439; p=1.480E-58) as well as the gene and the exon region (r=0.101; p=0.001). The higher correlation with the 3’ UTR region is to be expected, due to a higher abundance of probes designed in these regions.

160 genes are found to be differentially expressed using the gene-based approach only, with three not included in the region-based CDF. Further examination shows that 122 of these (78%) have a gene-based p-value > 0.03, and 80 (50%) have a gene-based p-value > 0.04, indicating the detected differences are just below the cutoff level. Analysis of the region-based p-values show that 120 of these (77%) have a region-based p-value < 0.10, and 146 (94%) have a region-based p-value < 0.20, putting these genes just above the significance threshold.

An additional 423 genes are found to be differentially expressed using the region-based approach only, with 203 from the 3’ UTR only, 10 from the 5’ UTR only, 206 from the exon only, and 4 from both the 3’ UTR and exon. Unlike the DEGs uniquely found in the gene-based approach, those genes found to be differentially expressed in the region-based approach typically have a much higher p-value in gene-based analysis, with only 31% having a p-value between 0.05 and 0.10. This supports our reasoning that separating into functional regions allows detection of subtle changes in transcript formation that may have a larger functional impact of those transcripts which has been further validated by experimental work showing differential expression of the 3’ UTR of the CAMKIV gene plays a role in localization [23].

In order to determine why some genes were only detected by brainarray, we examined probes of those genes. 39 probes were not used in our CDFs since they aligned to multiple locations in the rn6 genome. 10 probes did not match gene structures in Ensembl and were not used in the CDFs. 18 probes were removed because the probe set contained fewer than three probes. 40 probes were used in different probe sets other than those annotated by brainarray.

In the GSE48611 data series, DEGs were determined for two pairwise comparisons: isogenic Ts21 vs. control iPSCs for both DS1 and DS4. We reanalyzed the data using region, gene and the original Affymetrix^®^ supplied CDF obtained from the Affymetrix^®^ Netaffx™ web site. For DS1, our gene-based CDF identified an additional 194 DEGs not found using the original CDF and 616 DEGs identified by both methods. For DS2, our gene base CDF identified an additional 331 DEGs found using our method only and 337 DEGs identified by both methods (Table 8).

**Table 8.** DEGs detected by our gene based CDF and GPL570.

| Cases | Our Gene Based CDF | GPL570 | Common |
| --- | --- | --- | --- |
| DS1 versus Ts21 | 810 | 2421 | 616 |
| DS2 versus Ts21 | 668 | 1840 | 337 |

## Discussion

One of the limitations of microarray technologies is the design of probes based on available sequence and annotation data at the time of design. Based on our analysis, the percentage of uniquely mapping probes varies from 84% (rat) to 87% (human), indicating that changing knowledge about the genome itself plays a role in probe utilization. In terms of annotation, the rat genome is known to have more incomplete information when compared to mouse and human, which is reflected in the fact that only 47% of the rat probes lie in region-based locales (exons and UTRs) compared to 65% for mouse, and 69% for human. Since this can potentially lead to a small number of probes in each annotated region (and thus increased false positive rates), we have further required at least three probes be present in each probe set for our analysis. Both unrestricted (1 or more) and restricted (3 or more) probe groupings are available as CDFs. To further illustrate the importance of region-based CDFs, using the subset of 4,024 genes with probesets in both the CDS and 3’ UTR regions, we were able to identify 203 differential expression events at the 3’ UTR level that do not show differential expression within the CDS. In addition, these events are not detected using the standard Affymetrix^®^ CDF. Further analysis of these 203 genes yields some genes of particular interest. For instance, the 3’ UTR of *GRIK4* (Glutamate Ionotropic Receptor Kainate Type Subunit 4) was up-regulated (p-value 0.0450) while the CDS was not significantly regulated (FC=1.07; p-value 0.4525), suggesting the 3’ UTR of this gene was lengthened (Fig. 4). *GRIK4* regulates kainite-receptor signaling and neuroplasticity [39] and its missregulation is associated with neurological diseases including Alzheimer’s [40], bipolar disorder [41], and others. Interestingly, a deletion variant specific to the 3’ UTR of *GRIK4* is protective of bipolar disorder [41]. Alongside our observation, this suggests that regulation of this plasticity-associated gene occurs though its 3’ UTR. We also observed that the 3’ UTR of *VEGFA* (vascular endothelial growth factor-A) was downregulated (-1.17 FC; p=0.0102) and expression of its CDS was unchanged (1.01 FC; p=0.8334) (Fig. 5). The 3’UTR of *VEGFA*, a potent neuromodulator, undergoes a well-described binary switch to regulate its expression [42]. Our observations suggest the *VEGFA* 3’ UTR undergoes an additional layer of regulation by shortening during collateral sprouting.

**Fig. 4.**
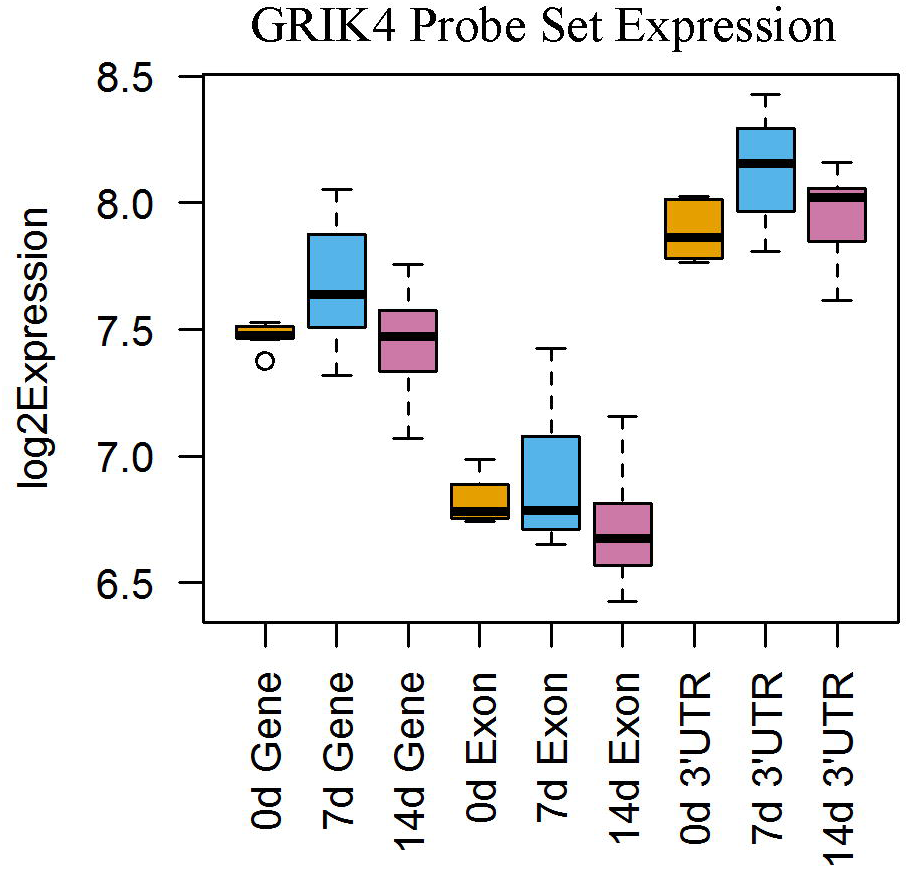
GRIK4 Probe set expression levels within the gene, exon, and 3’ UTR regions.

**Fig. 5.**
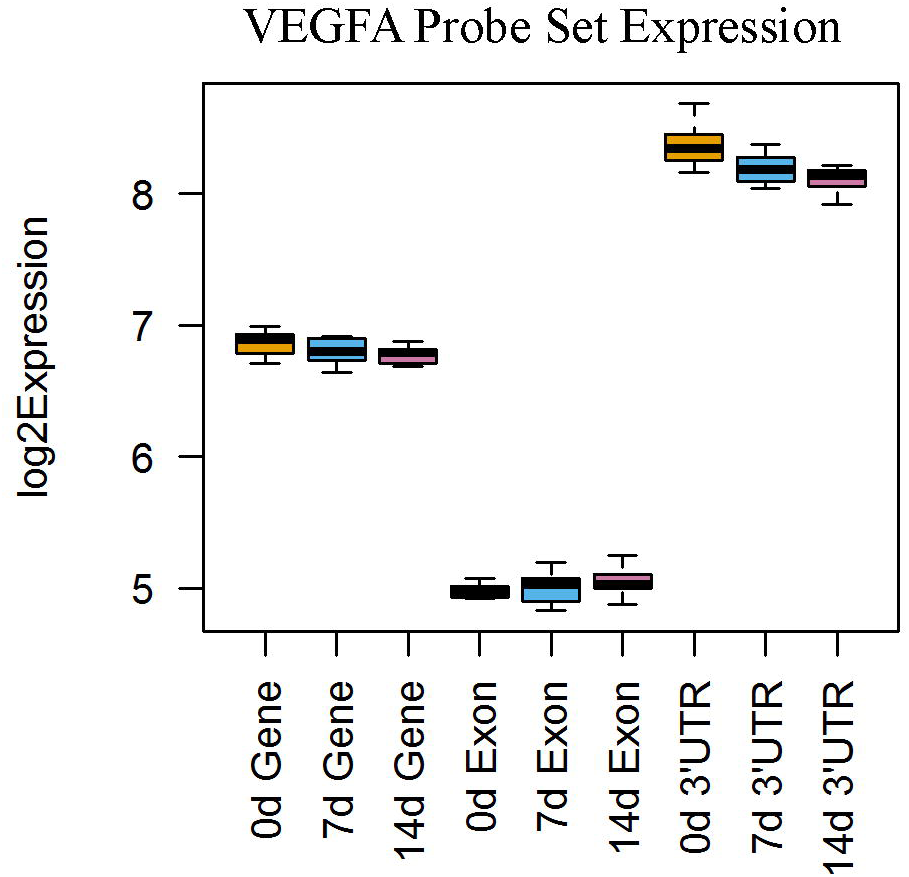
VEGFA Probe set expression levels within the gene, exon, and 3’ UTR regions.

As our analysis with the GSE48611 and GSE72551 datasets show, reanalysis of publicly available datasets using updated annotations can yield additional information when compared to the use of the original CDFs. In our case, the region-based CDFs allow for a better understanding of 3’ UTR dynamics through the reanalysis of publicly available data. While current high-throughput sequencing technologies may allow for a more complete picture, this custom CDF approach will allow for deeper insight with only minimal computational cost, taking advantage of the high volume of publicly available GeneChip^®^ data.

## Conclusions

We proposed a framework for reannotating and reassigning probe groups for Affymetrix^®^ GeneChip^®^ technology based on functional regions of interest. Our work differs from others in that we annotated probes in UTR and exon levels in addition to gene and transcript (isoform) levels. We illustrated how this framework affects the detection of differentially expressed genes, particularly when focusing on functional regions of interest. Removing probes that no longer align to the genome without mismatches or align to multiple locations can help to reduce false-positive differential expression, as can removal of probes in regions overlapping multiple genes.

The main motivations of our work was profiling the contribution of UTR and exon regions to the gene expression levels globally. Our results indicate that differentially expressed in either the gene-based or region-based CDF shows high concordance and separating out into functional regions allows for the detection of subtle changes in transcript formation.

## Acknowledgments

Support provided by National Institute of Health (NIH) grants P20GM103436 and R01NS094741. Its contents are solely the responsibility of the authors and do not represent the official views of the funding organization.

## Availability of data and materials

The datasets supporting the conclusions of this article are available in the figshare repository, http://10.6084/m9.figshare.3840144. Individual custom CDFs can also be accessed at: http://bioinformatics.louisville.edu/RegionCDFDesc.html

